# Disruption of LEDGF/p75-directed integration derepresses antisense transcription of the HIV-1 genome

**DOI:** 10.1101/2024.12.06.627169

**Authors:** Philip R. Tedbury, Darius Mahboubi, Maritza Puray-Chavez, Raven Shah, Obiaara B. Ukah, Claudia C. Wahoski, Hind J. Fadel, Eric M. Poeschla, Xinlin Gao, William M. McFadden, Maria Gaitanidou, Nikolaos Kesesidis, Karen A. Kirby, Thomas H. Vanderford, Mamuka Kvaratskhelia, Vasudevan Achuthan, Ryan T. Behrens, Alan N. Engelman, Stefan G. Sarafianos

## Abstract

Disruption of HIV-1 Integrase (IN) interactions with the host-factor Lens Epithelium-Derived Growth Factor (LEDGF)/p75 leads to decreased, random integration, increased latent infection, and described here, accumulation of HIV-1 antisense RNA (asRNA). asRNA increase was observed following interruptions of IN-LEDGF/p75 interactions either through pharmacologic perturbations of IN-LEDGF/p75 by treatment with allosteric HIV-1 integrase inhibitors (ALLINIs) or in cell lines with LEDGF genetic knockout. Additionally, by impairing Tat-dependent HIV transcription, asRNA abundance markedly increases. Illumina sequencing characterization of asRNA transcripts in primary T cells infected in the presence of ALLINIs showed that most initiate from within the HIV-1. Overall, loss of IN-LEDGF/p75 interactions increase asRNA abundance. Understanding the relationship between ALLINIs, integration sites, asRNA, and latency could aid in future therapeutic strategies.

## Introduction

Human immunodeficiency virus type-1 (HIV-1) replication requires dynamic interplay between viral and host factors. One critical step of HIV-1 replication is the integration of the viral genome into a host cell chromosome, which is catalyzed by HIV-1 integrase (IN) (*1-3*). Integration is not random, rather, it has been demonstrated to favor transcriptionally active and highly spliced gene-rich regions (*4-10*). Several cellular host proteins have been established to contribute to integration site distribution (ISD) by directly interacting with the pre-integration complex (PIC) (*11, 12*). Perturbation of proper ISD resulting from interference with these host proteins has been associated with viral latency (*11*). Latent HIV infection cannot be cleared by antiretrovirals currently available on the market and thus HIV latency is a significant barrier to an HIV-1 cure (*13, 14*).

Cleavage and polyadenylation specificity factor subunit 6 (CPSF6) is a host factor essential for effective HIV-1 infection (*15-19*); CPSF6 helps mediate ISD by binding to PIC-associated HIV-1 capsid protein (CA), promoting trafficking of PICs to highly active genes (*17*). Loss of CA-CPSF6 interaction leads to viral DNA integration into transcriptionally suppressed chromatin near the nuclear periphery (*18*). CPSF6 also plays a critical role in latency reversal and has been shown to promote viral transcription, highlighting its importance in HIV-1 transcriptional studies (*19*). Another host factor important for integration is LEDGF/p75, a transcriptional coactivator that is encoded by the PC4 And SRSF1 Interacting Protein 1 (*PSIP1*) gene. LEDGF is expressed as two isoforms, p75 and p52 (*20*). Both isoforms contain a chromatin binding PWWP motif and two AT-Hook motifs, which allow for direct interaction with the host chromosome (*10, 21-23*). LEDGF/p75 additionally possesses an integrase binding domain (IBD) that is absent in the p52 isoform (*24-26*). During HIV-1 infection, bimodal interactions of LEDGF/p75 with the PICs and host cell chromatin regulate the process of viral integration (*23, 27, 28*). These chromatin and DNA tethering domains are also critical in LEDGF endogenous activities, including its important roles in transcriptional regulation and modulating RNA splicing of stress-related genes, cell survival, and chromatin association (*21, 29-31*). The interaction between LEDGF and IN directs HIV-1 proviral integration into intron-dense regions of the genome and towards internal regions of genes (*17*). Allosteric IN inhibitors (ALLINIs), a.k.a. LEDGF-IN site inhibitors (LEDGINs) or non-catalytic site IN inhibitors (NCINIs), are a class of antiretroviral agents that bind at the LEDGF-binding site in IN abrogating binding of LEDGF/p75 (*24, 32-35*). ALLINIs also interrupt IN-RNA interactions *via* drug-induced IN aggregation, leading to the formation of aberrant virions (*36, 37*). Two members of the ALLINI family recently entered clinical trials (*35, 38*).

Perturbation of the interaction between IN and LEDGF promotes a) greater propensity for random integration, b) integration into sites closer to transcription start sites (TSSs) and CpG islands, and c) increases the frequency of HIV-1 latent infection (*17, 28, 39, 40*). Antisense transcription has been reported in retroviruses and has been shown to have both coding and noncoding roles: the Hbz transcript of HTLV-1, as one example, plays an important role in the HTLV-1 replication cycle and in the pathogenesis of adult T cell leukemia (*41, 42*).

Initial reports of an HIV-1 antisense RNA transcript (asRNA) (*43, 44*) were followed by *in vitro* studies reporting HIV-1 transcription of multiple antisense transcripts that were thought to originate from the HIV-1 3’ long terminal repeat (LTR) (*45, 46*). A subset of asRNA transcripts have been shown to encode a transmembrane protein, antisense protein (ASP) (*47*). People living with HIV-1 (PLWH) have been reported to produce antibodies against ASP (*48, 49*); however, the role of ASP in HIV-1 replication and viral pathogenesis remains unclear. HIV-1 asRNA exists as a mixed population of transcripts that can vary in length and degree of post-transcriptional modifications (*44, 45, 50, 51*). There are HIV-1 asRNAs described to function similarly to long non-coding RNA (lncRNA) to regulate the transcription of active genes (*45, 52*). One model proposes that HIV-1 asRNA may recruit the polycomb repressive complex 2 (PRC2) to transcriptionally repress HIV-1 replication and promote viral latency (*53*). Nonetheless, the mechanism by which transcription of HIV-1 asRNA is affected by viral and cellular factors remains unclear.

To date, LEDGF/p75 and HIV-1 asRNA have been separately linked to HIV-1 latency (*39, 46, 53*). Here we report a direct relationship between the loss of IN-LEDGF interactions and increased asRNA levels, which suggest an interlinked mechanism of promoting viral latency. We identified this relationship in various cell types by employing MICDDRP (multiplex immunofluorescent cell-based detection of DNA, RNA and Protein), which is based on RNAscope^TM^ branched *in situ* hybridization technology combined with immunofluorescence for visualization of viral RNA (vRNA), nuclear DNA, and Gag simultaneously (*54*). To validate the relationship between HIV-1 asRNA and loss of IN-LEDGF/p75, we used *PSIP1* knock-out (*PSIP1* -/-) cells and ALLINIs to circumvent the IN-LEDGF interactions during HIV-1 integration. Finally, we characterized viral genome-wide plus-sense and asRNA transcripts by Illumina sequencing to inform the mechanisms underlying asRNA production and its possible role in HIV-1 replication.

## Results

### Loss of IN-LEDGF/p75 interactions increases frequency of cells expressing HIV-1 asRNA

LEDGF/p75 and CPSF6 have both been reported to play distinct roles in integration and subsequent HIV-1 transcription. Therefore, HIV-1 transcription was examined in cells lacking either or both of these cellular proteins. We measured the proportion of cells expressing HIV-1 sense RNA ((+)RNA), antisense RNA ((-)RNA), or both ((+/-)RNA) in HEK293T LEDGF knock-out cells (LKO), CPSF6 knock-out cells (CKO), and LEDGF/CPSF6 double knock-out (DKO) cells (*17*). Cells were fixed and labeled for viral RNA (vRNA) using *in situ* hybridization and images were captured using a BioTek Cytation 5 Cell Imaging Reader.

The HEK293T LKO and DKO cells revealed 4- and 11-fold increases in total asRNA expressing cells ((-)RNA and (+/-)RNA cells), respectively, when compared to isogenic-matched wild-type (WT) cells. By contrast, HEK293T CKO cells did not exhibit a significant change in the proportion of cells expressing asRNA (Fig. 1A). Experiments performed in the Jurkat T-cell line revealed similar results, with incidence of asRNA expressing cells increasing by 5-fold in LKO cells relative to WT (Fig. 1B).

**Fig. 1.**
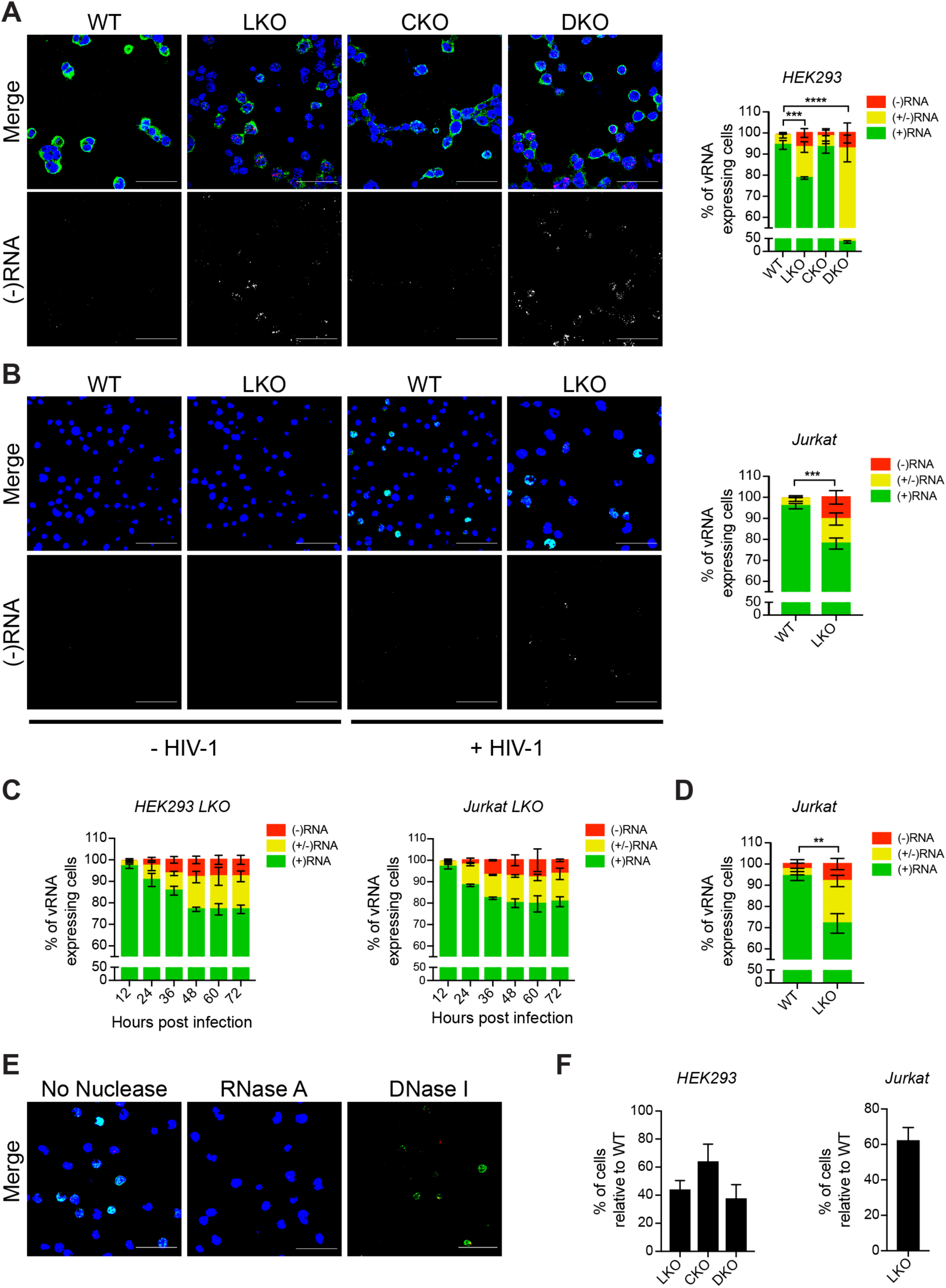
FIG 1 Levels of asRNA in CPSF6 and LEDGF knock-out cell lines. (**A**) HEK293T cells lacking *PSIP1* (LEDGF-/-, LKO), CPSF6 (CKO), or both (DKO) were infected with VSV-G-pseudotyped HIV-1 at a multiplicity of infection (MOI) of 1, fixed 48 hpi, and sense RNA (green), antisense RNA (red), and nuclei (blue) were stained. Cells expressing either sense RNA only ((+)RNA), antisense RNA only ((-)RNA), or both ((+/-)RNA) were counted using Gen5 software and expressed as percentages. The lower row highlights (-)RNA signals in the absence of red color. (**B**) WT and LKO Jurkat cells were infected and scored as described in (A). (**C**) Time course of (+)RNA, (-)RNA, and (+/-)RNA expression. LKO HEK293T and Jurkat cells were infected at an MOI of 0.5 and samples were fixed every 12 hpi, for up to 72 h. (**D**) Proportion of cells expressing vRNA for WT and LKO Jurkat cells was determined by flow cytometry. (**E**) HIV-1 infected Jurkat cells were fixed then treated with vehicle, RNase A, or DNase I prior to nucleic acid probing. (**F**) Infection of HEK293T- and Jurkat-derived knockout cell lines relative to WT is reported using sense vRNA as the indicator of infection and measured using a cell imaging multimode reader. Number of infected cells in knock-out lines is plotted as a percentage of WT. For all scoring conditions, *n* = 3 independent experiments. At least 5 fields of view containing a total of >500 vRNA expressing cells were scored for each experiment, with standard deviations shown. Scale bars represent 80 µm. ** p ≤ 0.01, *** p ≤ 0.001, **** p ≤ 0.0001 was determined by Tukey’s multiple comparison test for all cells expressing antisense RNA (i.e. (-)RNA and (+/-)RNA cells).

To determine the rate of asRNA production, we measured HIV-1 asRNA-positive cells in LKO HEK293T and Jurkat cells at 12-h intervals for 72 h, revealing that transcript levels peaked at 48 h post-infection (hpi) and remained at that level until at least 72 hpi; therefore, all asRNA levels were monitored and quantified using data from the 48-h timepoint for subsequent experiments (Fig. 1C).

To validate the scoring method for these studies, we independently quantified the asRNA-positive Jurkat LKO cells versus WT by flow cytometry, which also yielded a 5-fold increase in cells expressing asRNA in LKO cells (Fig. 1D). To confirm that the *in situ* hybridization was labeling vRNA, a series of nuclease treatments with DNase I and RNase A were performed (Fig. 1E). RNase A degraded the sense and antisense vRNA fluorescent signals while DNase I only depleted the nuclear DNA signal. Finally, knock-out cell lines exhibited reduced frequency of infected cells relative to WT, consistent with the roles of CPSF6 and LEDGF/p75 as cofactors in nuclear import and integration (Fig. 1F).

As an additional control, we used the complementary approach of perturbing IN-LEDGF interactions pharmacologically, to verify that changes in asRNA levels derived from the loss of the IN-LEDGF interactions and not from indirect effects on host physiology imparted by the loss of LEDGF proteins. We treated HEK293T (Fig. 2A), Jurkat (Fig. 2B), and primary CD4^+^T cells (Fig. 2C) with the ALLINIs BI-D and BI-224436 (*3, 55*), which disrupt IN-LEDGF/p75 interactions by binding to the LEDGF/p75-binding pocket on IN to compete with LEDGF/p75 binding. ALLINIs are largely non-toxic to cells and should thus have little or no impact on the role of LEDGF/p75 in host biology. HEK293T and Jurkat cells exhibited 4.5-fold and 5.5-fold increases, respectively, in the proportion of asRNA-positive cells when treated with BI-D compared to the DMSO control (Fig. 2A and B). These values are similar to the changes observed in the LEDGF/p75 knock-out cell lines. Primary CD4^+^T cells treated with BI-D similarly revealed a 3.6-fold increase in the proportion of asRNA-positive cells (Fig. 2C).

**Fig. 2.**
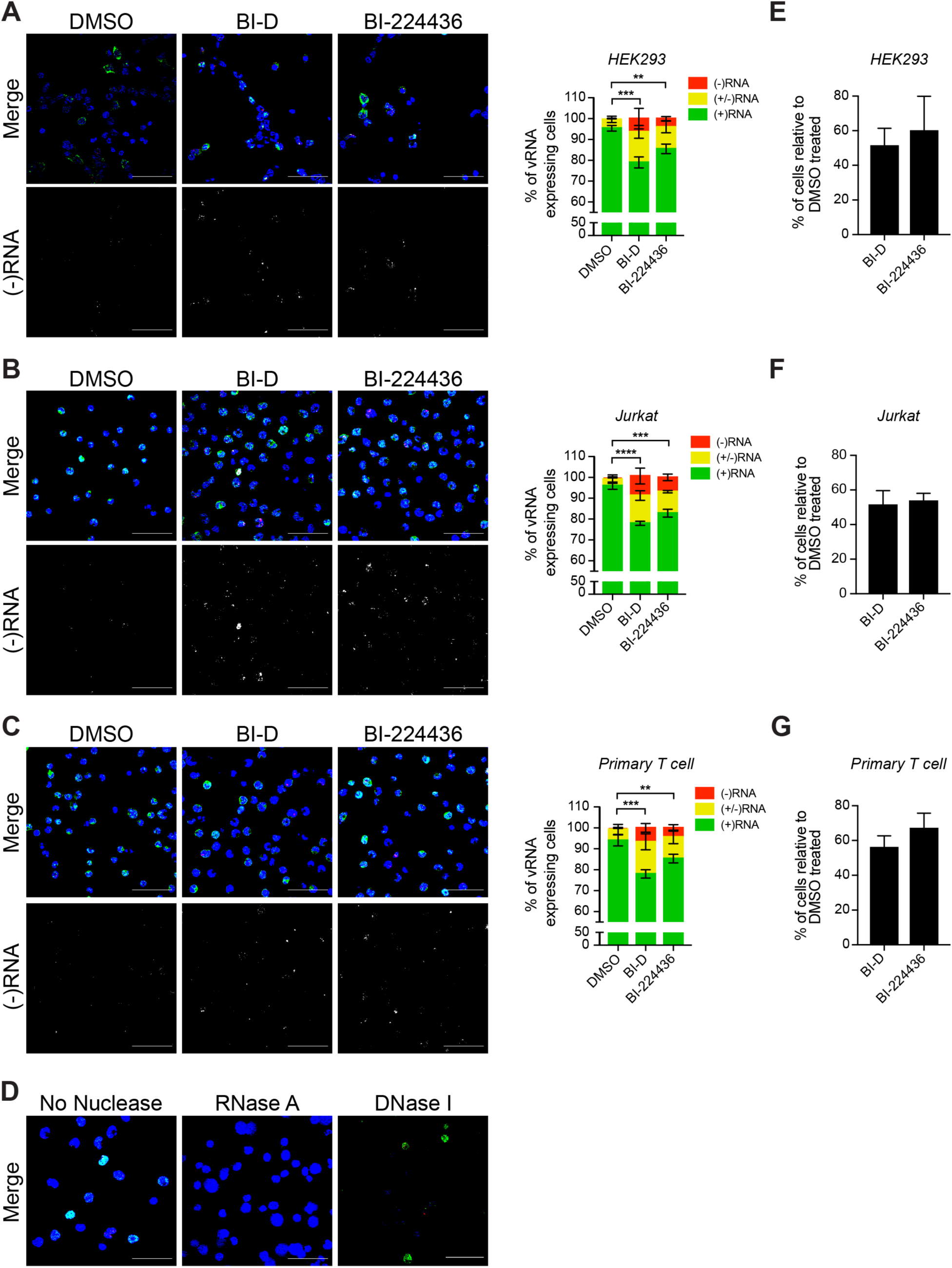
Levels of asRNA following inhibition of the IN-LEDGF interaction following ALLINI treatment. (**A**) HEK293T, (**B**) Jurkat, and (**C**) primary CD4^+^T cells were treated with DMSO or 10 μM of BI-D or BI-224436 12 h prior to infection with HIV-1 at an MOI of 1. Cells were fixed 48 hpi and then sense RNA (green), asRNA (red), and nuclei (blue) were stained. Cells expressing sense RNA ((+)RNA), asRNA ((-)RNA) or both vRNAs ((+/-)RNA) were counted using Gen5 software and plotted as percentages of total infected cells. (**D**) HIV-1 infected Jurkat cells were fixed then treated with vehicle, RNase A, or DNase I prior to nucleic acid probing. (**E-G**) Infection of ALLINI-treated cells was quantified by counting the number of cells expressing vRNA compared to the total number of cells in all the fields of view. The percentage of infected ALLINI-treated cells was plotted relative to DMSO-treated cells. For all scoring conditions *n* = 3 independent experiments. At least 5 fields of view containing a total of >500 vRNA expressing cells scored for each experiment, with standard deviations shown. Scale bars represent 80 µm. ** p ≤ 0.01, *** p ≤ 0.001, **** p ≤ 0.0001 was determined by Tukey’s multiple comparison test for all cells expressing antisense RNA (i.e. (-)RNA and (+/-)RNA cells).

Similar data were acquired using BI-224436; the increases in the proportion of asRNA-positive cells were 4-fold (HEK293T), 3.2-fold (Jurkat), and 2.2-fold (primary CD4^+^ T cells) (Fig. 2A-C). Again, nuclease treatment controls were used to confirm the identity of the labeled nucleic acid as RNA (Fig. 2D). We also measured viral infectivity in ALLINI-treated cells and found that BI-D and BI-224436 decreased HIV-1 infectivity by approximately 50% relative to DMSO-treated control (Fig. 2E-G). Together, these studies show a consistent trend among different cells and treatments (genetic knock-out versus pharmacological inhibition) indicating that asRNA incidence increases relative to total vRNA expressing cells when IN-LEDGF/p75 interactions are impaired.

### Disruption of IN-LEDGF/p75 interactions leads to increased asRNA expression and decreased sense RNA levels

As the perturbation of the IN-LEDGF/p75 complex during integration resulted in suppressed HIV-1 infection and increased HIV-1 asRNA-positive cells, we continued by assessing the levels of sense and antisense RNA in HIV-1 infected cells. The probes used in RNAscope^TM^ experiments were different for sense and asRNA, so the absolute amounts between these two species could not be compared. Amounts of sense and asRNA were determined relative to WT or DMSO-treated controls, allowing us to assess the effects of disrupting the IN-LEDGF/p75 interaction. Average fluorescence intensity of sense (Fig. 3A, *left*) and antisense (Fig. 3A, *right*) RNA were quantified in HIV-1 infected WT and LKO Jurkat cells. Fluorescence intensity of sense RNA was reduced by 36% in the LKO Jurkat cells relative to WT. In contrast, the fluorescence intensity for asRNA increased by 104% in the LKO cells (Fig. 3A). Data acquired in HEK293T-derived cells were comparable to those in Jurkat cells, with sense RNA reducing by 67% (fig. S1A, *left*) and asRNA increasing by 247% (fig. S1A, *right*) in LKO cells compared to their WT counterparts.

**Fig. 3.**
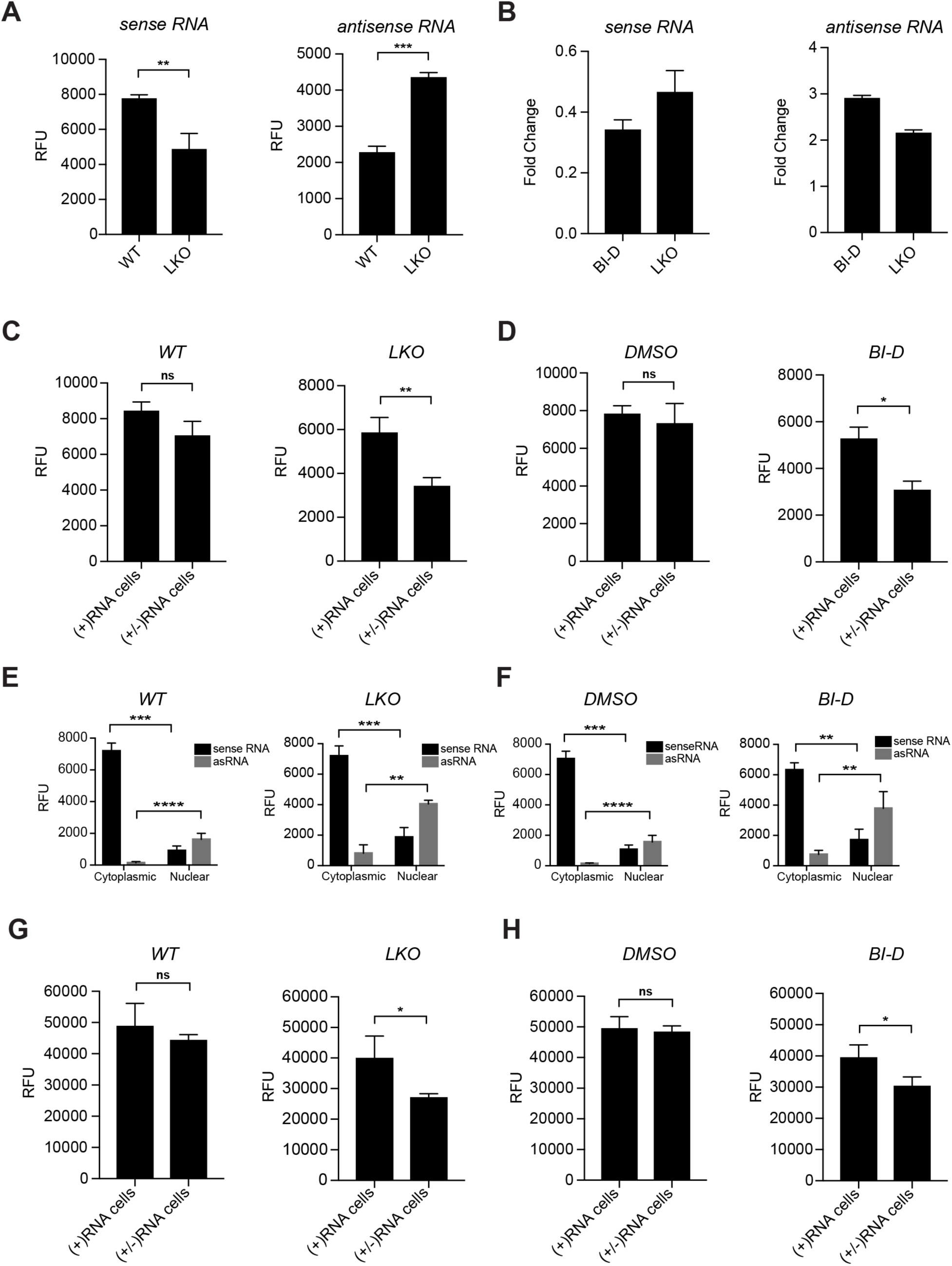
vRNA quantity and localization following the loss of IN-LEDGF/p75 interaction. Jurkat cells were infected with HIV-1 at an MOI of 0.4, fixed 48 hpi and stained for sense RNA and antisense RNA. Images were captured using a BioTek Cytation 5 multimode reader and fluorescence intensity was measured using Gen5 software. (**A**) Fluorescence intensity of sense RNA (*left*) and asRNA (*right*) in WT and LKOJurkat cells. (**B**) HIV-1 sense RNA (*left*) and asRNA (*right*) levels in LKO and BI-D-treated cells relative to WT and DMSO-treated cells, respectively, determined by RT-qPCR. (**C**) HIV-1 sense RNA levels in WT (*left*) and LKO (*right*) Jurkat cells containing either exclusively sense RNA ((+)RNA) or both vRNA transcripts ((+/-)RNA). (**D**) Mean fluorescence intensity of sense RNA in DMSO (*left*) or BI-D-treated (*right*) Jurkat cells in (+)RNA or (+/-)RNA expressing cells. (**E** and **F**) Mean fluorescence intensity and localization of vRNA in individual Jurkat cells was measured using Gen5 software. Hoechst 33342 and high content screening (HCS) CellMask™ Deep Red to segment nuclear and cytoplasmic compartments, respectively. (**G** and **H**) p24 abundance in (+)RNA and (+/-)RNA expressing Jurkat cells, as determined by mean fluorescence intensity following immunofluorescent labelling. For all scoring conditions *n* = 3 independent experiments. At least 5 fields of view containing a total of >500 vRNA expressing cells were scored for each experiment, with standard deviations shown. * p ≤ 0.05, ** p ≤ 0.01, *** p ≤ 0.001, ns, not significant.

To further validate the fluorescence intensity measurements, we used RT-qPCR to compare relative levels of sense and antisense RNAs. Sense RNA levels showed a 2.8- and 2.4-fold reduction in BI-D-treated and LKO samples compared to DMSO-treated and WT controls, respectively (Fig. 3B, *left*). Consistent with our fluorescence intensity data, RT-qPCR analysis of asRNA levels exhibited 2.9- and 2.1-fold increases for BI-D-treated and LKO Jurkat cells, respectively (Fig. 3B, *right*). In summary, both RNAscope^TM^ and RT-qPCR showed that when the IN-LEDGF interaction was lost, sense RNA levels were reduced, whereas asRNA levels were increased.

Having demonstrated a correlation between the loss of IN-LEDGF interactions and the amount of sense and asRNA at the level of bulk cell population, we proceeded to examine whether the presence of asRNA transcripts in cells containing both vRNAs was associated with changes in sense RNA levels, compared to cells exclusively expressing sense RNA. The fluorescence intensity of sense RNA was not significantly reduced in the presence of asRNA in WT Jurkat cells, however, in LKO cells, sense RNA levels decreased by 34% in the presence of asRNA (Fig. 3C). Similar data were observed in HEK293T cells (fig. S1B). These results indicate a link between expression of high levels of asRNA seen in the LKO cells and a reduction in levels of sense RNA. We verified this relationship in ALLINI-treated cells, which produced similar outcomes to the LKO cells; DMSO-treated Jurkat cells did not display a significant reduction of sense RNA in the presence of asRNA, but the BI-D-treated cells exhibited a 26% loss in sense RNA in the presence of asRNA (Fig. 3D). Again, similar data were obtained with ALLINI-treatment experiments with HEK293T cells (fig. S1C).

### Localization of HIV-1 sense and antisense RNAs and p24 expression

As the function of viral RNA depends not only on quantity but also on its location, we measured nuclear and cytoplasmic levels of HIV-1 asRNA in LKO and ALLINI-treated cells. Antisense RNA was almost exclusively localized to the nucleus in all cells in which it was found, while HIV-1 sense RNA was found primarily in the cytoplasm (Fig. 3E and F).

The decrease in HIV-1 sense RNA levels in the presence of asRNA could be indicative of viral replication suppression by modulating gene expression. To determine if the loss of sense RNA levels in the presence of asRNA led to a decrease in HIV-1 capsid protein (CA or p24), we used MICDDRP for simultaneous visualization of vRNA, nuclear DNA, and Gag (*54*). Fluorescence intensity from p24 labeling in WT and LKO Jurkat cells was measured to investigate the association between the presence of asRNA and CA levels. The average CA levels only decreased by 36% in the presence of asRNA produced from LKO Jurkat cells (Fig. 3G, *right*) and did not decrease in wild-type cells (Fig. 3G, *left*). LKO HEK293T cells also showed an average decrease in CA levels of 30%, while no change was observed in the WT cells (fig. S1D). CA signal also decreased by 33% in (+/-)RNA cells for BI-D-treated samples (Fig. 3H, *right*) when compared to (+)RNA cells but was unchanged in DMSO-treated cells (Fig. 3H, *left*), suggesting that only asRNA transcript(s) produced when IN-LEDGF/p75 interactions were lost affects capsid protein levels. These data were recapitulated in BI-D-treated HEK293T cells, which exhibited a loss of average CA signal by 31% in (+/-)RNA containing cells, while no difference was measurable between (+)RNA and (+/-)RNA cells when treated with DMSO (fig. S1E).

Hepatoma-derived growth factor-related protein 2 (HDGFL2), which also contains IBD and PWWP domains (*26*), is a paralog to LEDGF/p75 and has been reported to direct ISD in the absence of LEDGF/p75 (*56, 57*). To determine if the loss of both IBD-containing proteins promotes greater asRNA production, we quantified the number of cells expressing HIV-1 sense RNA (+RNA), asRNA (-RNA), or both (+/-RNA) in WT, LKO, HDGFL2 knock-out (HKO), and LEDGF/ HDGFL2 double knock-out (LHKO) Jurkat cells. HDGFL2 knock-out alone did not induce any changes in the proportion of cells expressing asRNA, but LHKO cells had increased asRNA production by 116% when compared to LKO cells (fig. S2A). The susceptibility of the Jurkat-derived knock-out cells to infection was quantified relative to the WT control cells. LHKO cells supported significantly less infection, similar to LKO cells (fig. S2B). Because LHKO increased the number of asRNA producing cells, we also quantified the amount of sense or antisense RNA to determine whether the levels of vRNA also changed. Depletion of both LEDGF/p75 and HDGFL2 did not result in any changes to the mean fluorescence intensity for HIV-1 sense RNA when compared to LKO cells (fig. S2C, *left)*. Likewise, quantification of average fluorescence intensity for labeled HIV-1 asRNA revealed that LHKO cells did not have increased asRNA levels relative to LKO cells (fig. S2C, *right*). Thus, similar to our conclusions on the CA-CPSF6 interaction, the IN-HDGFL2 interaction does not seem to contribute to the regulation of HIV-1 asRNA production.

### Disruption of 5’ LTR-mediated transcription leads to increased asRNA levels

The trans-activator of HIV-1 (Tat) is essential for viral transcription. Tat binds to the viral transactivation response (TAR) element in the 5’ untranslated region of HIV-1 RNA and recruits P-TEFb to phosphorylate RNA polymerase II complexes and promote transcription elongation (*58-60*). To begin to understand the origin of HIV-1 asRNA, we first asked whether transcription of HIV-1 asRNA is Tat-dependent. Two Tat-deficient viruses were used; one virus contains a stop codon in place of the Tat start codon, which prevents expression of Tat (*61*). The second Tat-deficient virus has a single base-pair substitution in the cysteine-rich domain creating a missense mutation and abolishing the trans-activation activity (*62, 63*). Typically, vRNA or HIV-1 capsid protein staining are used to measure readouts for HIV-1 infectivity; however, because neither of these methods are suitable for transcription-deficient viruses, we used Alu qPCR measurement of integrated proviruses to standardize virus stocks (Fig. 4B). Jurkat cells infected with Tat-deficient virus were fixed and stained for sense and asRNA. As expected, in the absence of functional Tat, no HIV-1 sense RNA was seen. To confirm that the Tat-deficient viruses were replication competent, except for the lack of Tat, we rescued HIV-1 sense RNA transcription by transfecting a Tat-expression vector into cells prior to infection (Fig. 4A). Quantitation of asRNA revealed that not only was asRNA synthesized in the absence of Tat activity, but expression was increased 4-fold compared to cells infected with WT HIV-1 (Fig. 4C). The proportions of cells expressing sense RNA, asRNA, or both transcripts were quantified, revealing that >95% of cells infected with Tat-deficient virus expressed asRNA; the WT distribution was restored when Tat was reintroduced by transfection with an expression vector (Fig. 4D).

**Fig. 4.**
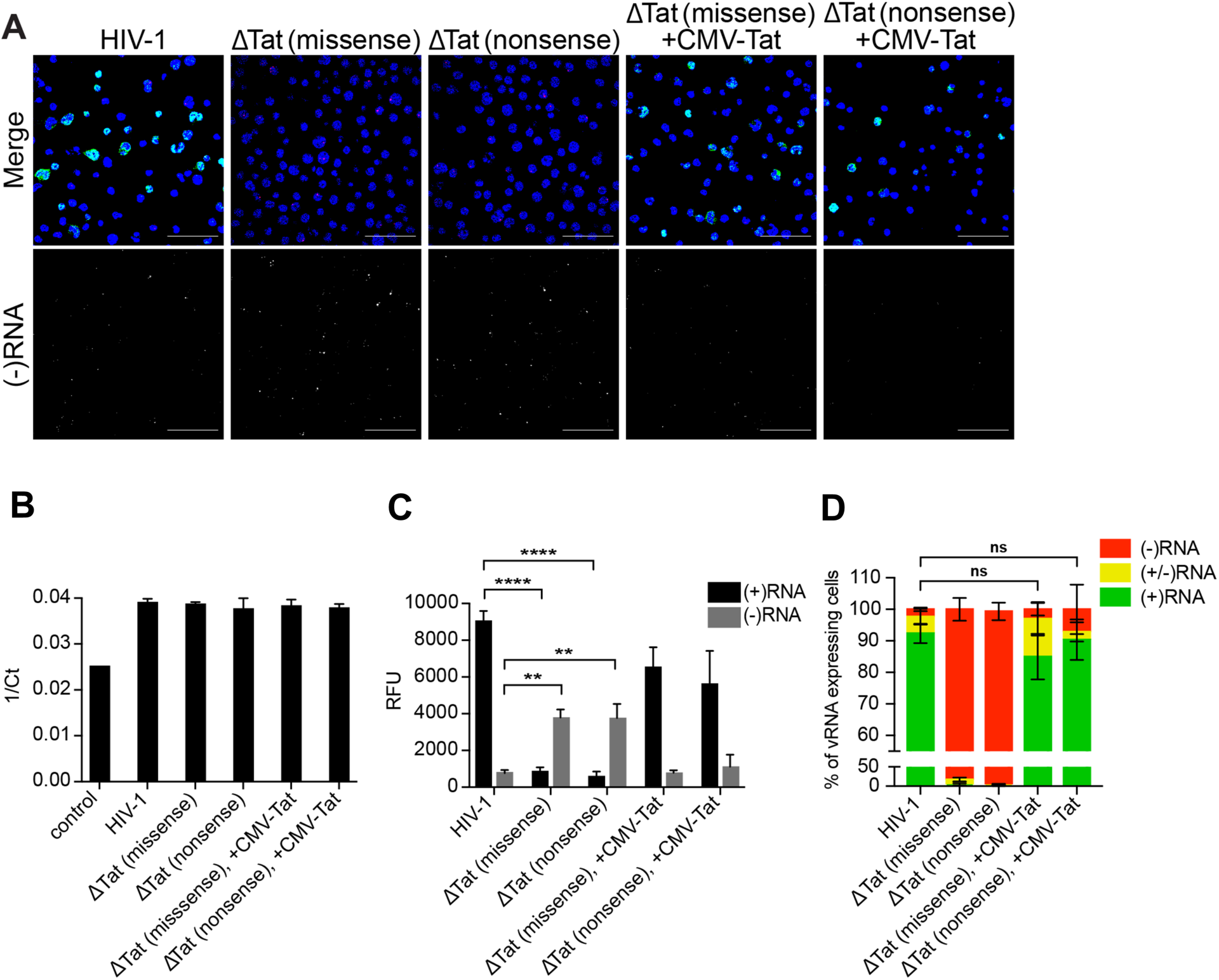
vRNA quantity and localization following the loss of IN-LEDGF/p75 interaction. Antisense RNA levels following infection with Tat-deficient virus. (**A**) Jurkat cells were infected with Tat-deficient HIV-1 virus and probed for sense (green) and antisense RNA (red) 48 hpi. Two Tat mutants were used for this study, one with a missense mutation and the other with an early nonsense mutation. A Tat expression vector was co-transfected to rescue (+)RNA expression of the Tat-deficient mutants. (**B**) Infection of target cells was confirmed and standardized using Alu qPCR. (**C**) Mean fluorescence intensity of sense and antisense RNA in actively transcribing cells infected with Tat-deficient HIV-1 following MICDDRP labeling and imaging on a BioTek Cytation 5 multimode reader was measured using Gen5 software. (**D**) Proportions of cells expressing exclusively sense RNA ((+)RNA), antisense RNA ((-)RNA), or both vRNAs ((+/-)RNA) were quantified for cells infected with Tat-deficient virus and plotted as percentages. For all scoring conditions *n* = 3 independent experiments. At least 4 fields of view containing a total of >300 vRNA expressing cells were scored for each experiment, with standard deviations shown. Scale bars represent 80 µm. ** p ≤ 0.01, **** p ≤ 0.0001 was determined by Tukey’s multiple comparison test. ns, not significant.

The data indicated that Tat was not required for expression of HIV-1 asRNA. Instead, asRNA levels appeared significantly elevated in the absence of Tat. To validate this observation, we employed an alternative model: Jurkat cells were engineered to express P-TEFb mutants that severely attenuate Tat-dependent transcription. WT, LKO, and P-TEFb mutant Jurkat cell lines were infected with HIV-1 and Alu qPCR analysis was used to quantify integrated vDNA and confirm consistent infection across the cell types (Fig. 5A and B). Compared to WT Jurkat cells, asRNA fluorescence intensity exhibited approximately 4.5-fold increase in the two P-TEFb mutant cell lines (Fig. 5A and C). We also observed a loss of sense RNA, presumably as a consequence of the inability of the HIV-1 Tat to recruit the mutant P-TEFb to RNA transcription complexes. The percentage of cells expressing asRNA (i.e. (-)RNA and (+/-)RNA) increased by approximately 10-fold for both mutant cells compared to WT cells (Fig. 5D). These data show that asRNA expression does not require HIV-1 Tat and is increased in the absence of trans-activator activity.

**Fig. 5.**
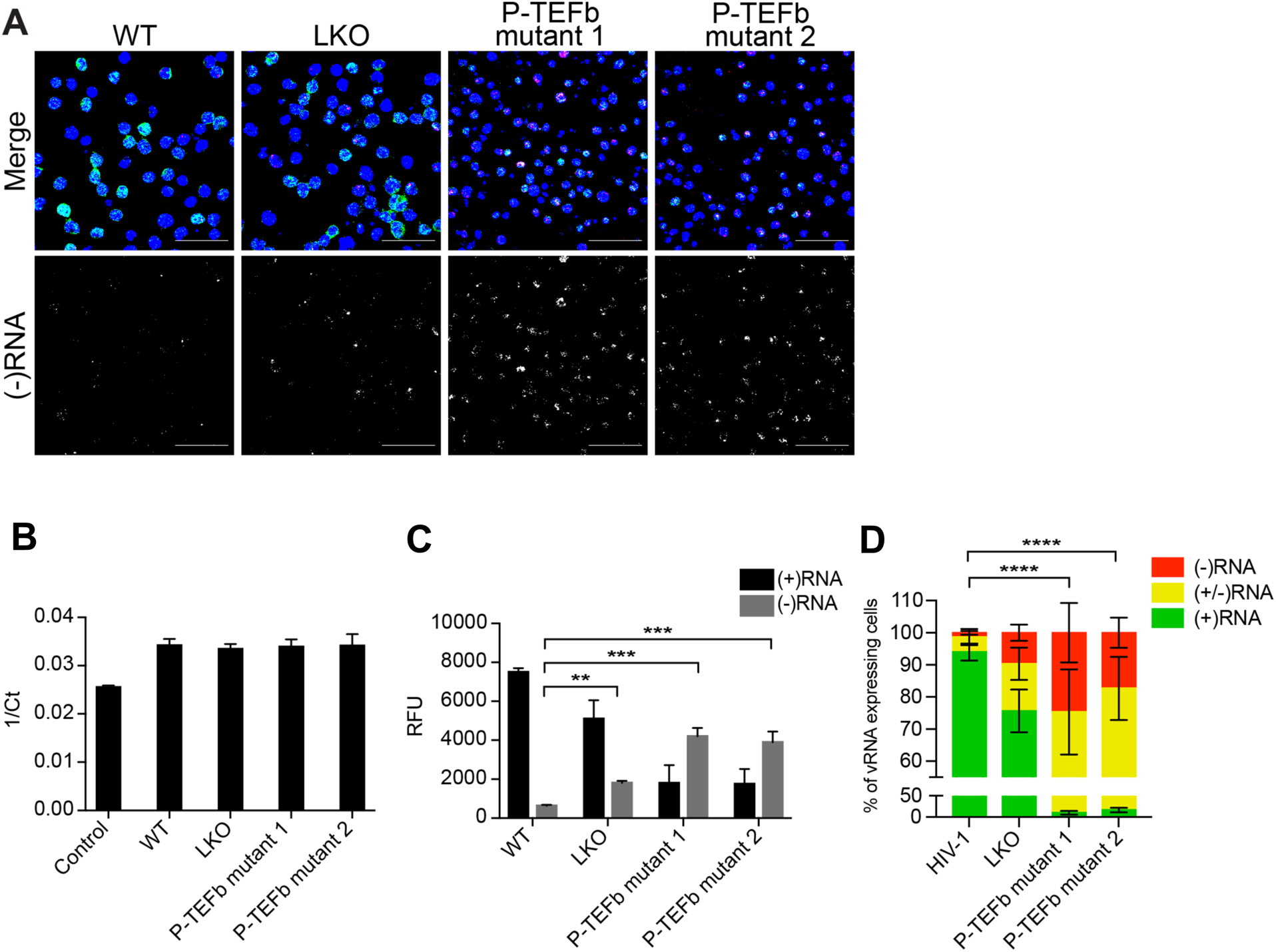
FIG 5 Antisense RNA levels in P-TEFb mutant cell lines. (**A**) WT, LKO and two P-TEFb mutant Jurkat cell lines were infected with HIV-1, fixed 48 hpi, and stained for sense RNA (red), antisense RNA (green), and nuclei (blue). (**B**) Alu qPCR was used to confirm proviral DNA levels. (**C**) Mean vRNA fluorescence intensity of infected cells. (**D**) Proportion of cells expressing exclusively sense RNA ((+)RNA), asRNA ((-)RNA), or both vRNA transcripts ((+/-)RNA) were quantified and plotted as percentage of total vRNA expressing cells. For all scoring conditions *n* = 3 independent experiments. At least 4 fields of view containing a total of >300 vRNA expressing cells were scored for each experiment, with standard deviations shown. Scale bars represent 80 µm. ** p ≤ 0.01, *** p ≤ 0.001, **** p ≤ 0.0001 was determined by Tukey’s multiple comparison test.

### Characterization of HIV-1 asRNA by Next Generation Sequencing (NGS)

The HIV-1 asRNA transcriptome is composed of transcripts varying in length, TSS, post-transcriptional modifications, and function (*44, 45, 50, 51*); therefore, we characterized the asRNA transcripts produced in the absence of the IN-LEDGF/p75 interaction. We performed Illumina HiSeq on RNAs isolated from HIV-1 infected human primary CD4^+^T cells treated with DMSO or BI-D (fig. S3 and S4A). Representative plots of read counts for HIV-1 sense and antisense RNA were generated for each treatment condition and mapped to the NL4-3 genome (fig. S3 and S4A). These data showed only minor changes in the distribution of HIV-1 sense transcripts, suggesting that the main effect on sense transcripts is a general reduction consistent with overall reduced infection levels under these conditions. However, we saw distinct changes in the pattern of asRNA reads.

To better visualize the changes in the respective vRNA read counts between DMSO- and ALLINI-treated samples, the reads at each position were expressed as the proportion of all the HIV-mapped reads. The difference between ALLINI reads and DMSO reads were plotted to show regions with the largest changes (fig. S4B). A region at ∼5,600 bp was found to be suppressed in the ALLINI-treated samples; the Alternative Splice Site Predictor (ASSP) (*64*) identified constitutive acceptor and donor splice sites at 5,965 bp and 5,708 bp, respectively, which suggested that the HIV-1 asRNA transcripts were spliced, consistent with a previous report (*51*), and that asRNA splicing was perturbed when transcribed from proviruses integrated in the presence of ALLINIs. A peak was also seen at the 3’ LTR at around 9,200 bp in a region overlapping *nef*, consistent with increased transcription in the vicinity of a proposed antisense TSS in the nef/3’LTR region (*50, 65*).

## Discussion

LEDGF/p75 binds directly to HIV-1 IN and tethers the PICs to chromatin during integration, favoring the mid-body of highly spliced genes (*17, 23, 24, 29, 57, 66*). Previous studies have separately revealed associations between the loss of IN-LEDGF/p75 interactions or HIV-1 asRNA, and latency (*39, 40, 45, 46, 51, 53*). In this study, we found that perturbation of the IN-LEDGF/p75 interaction during HIV-1 integration led to increased abundance of HIV-1 asRNA. Additionally, we used imaging and sequencing approaches to strengthen prior evidence that HIV-1 asRNA is a mixed population of transcripts whose abundance is inversely correlated with that of HIV-1 sense RNA, suggesting influence on HIV-1 gene expression and providing a potential mechanism whereby ALLINIs might impact not only HIV-1 assembly and maturation, and HIV-1 integration, but also expression from HIV-1 proviruses integrated in the presence of ALLINIs.

Various mechanisms could conceivably lead to HIV-1 asRNA production, including auto-integration, transcription from an antisense promoter in the HIV-1 3’-LTR or transcription from a host promoter, following integration of the HIV-1 provirus in the opposite sense to the host gene. Autointegration involves the integration of the LTR into the vDNA, potentially forming DNAs with alternating sense and antisense regions (*67*), and LEDGF/p75 has been proposed to influence autointegration (*68*). In this model, transcription would still be driven from the HIV-1 5’-LTR and as such would still be Tat dependent (*69*). Our data, however, indicated that production of asRNA was not dependent on Tat. In fact, asRNA levels were enhanced in the absence of Tat-mediated recruitment of P-TEFb, indicating transcription of HIV-1 asRNA does not originate from the 5’-LTR and that autointegration is probably not the mechanism underlying the production of asRNA in these experiments. This finding is supported by a previous study, which showed that co-expression of Tat and asRNA expression vectors led to a decrease of asRNA transcription *in vitro* (*44*). Additionally, deletion of the 5’-LTR has been shown to increase asRNA levels (*50, 70*).

To further characterize HIV-1 asRNA and better understand its transcription start site (TSSs), we employed Illumina NGS. HIV-1 asRNA is reported to be a mixed population of vRNAs, with little understood about the mechanisms that regulate expression, TSS, and post-transcriptional modifications. Most studies characterizing the asRNA transcripts have identified 3’-end polyadenylation (*44, 48, 50*), although one paper found the transcripts to lack a polyA tail (*45*). In addition, multiple TSSs have been reported to be in the U3 region, *nef* and *env* genes (*50, 65*), and transcription of HIV-1 asRNA can also originate from promoters in the host genome (*71-73*). Nevertheless, host-virus chimeras generally make up a small proportion of the total reads, and chimeras where the viral sequence is antisense make up a small subset of chimeric transcripts (*74, 75*). Our data supplement earlier findings, showing that although inhibiting IN-LEDGF/p75 interactions during infection suppressed the overall level of HIV-1 sense transcription, the pattern of sense RNA transcription was not appreciably altered. By contrast, the antisense transcripts exhibited distinct changes, most notably at the 3’ LTR, consistent with enhanced transcription from the proposed antisense TSS (*50, 65*), and in the center of the genome suggesting altered splicing of asRNA transcripts produced following integration in the absence of IN-LEDGF/p75 interactions. Although we observed chimeric HIV-1 antisense transcripts consistent with transcription originating in a host gene (not shown), the read numbers were too low to allow conclusions to be drawn about the potential role of IN-LEDGF/p75 interactions and the transcription of host-to-virus chimeric RNAs.

Given the potential for the asRNA to interact with both the proviral DNA and the sense RNA, we characterized sense RNA in cells expressing sense RNA only and those expressing both vRNAs. We found that only when IN-LEDGF/p75 interactions are perturbed, did the amounts of sense RNA and HIV-1 Gag protein decrease in the presence of asRNA. It is not clear why the reductions in sense RNA and protein only reached significance when looking at cells that both contained asRNA and lacked the IN-LEDGF interaction; it may be relevant that although asRNA-positive cells could be observed in both WT and LKO populations, the levels of asRNA in asRNA-positive LKO cells were higher than in WT cells. The increase in asRNA levels sequestered to the nucleus could function to suppress HIV-1 gene expression and replication, pushing these cells into self-directed state of viral latency with reduced reactivation potential as previously reported (*39*). However, we cannot establish a causal relationship from our data.

The associations between LEDGF/p75, asRNA, and HIV-1 latency merit further investigation, particularly given the recent introduction of the ALLINI STP040, also known as pirmitegravir (NCT05869643) (*38, 76*), into clinical trials. The association with latency could have either an adverse effect on HIV cure efforts, by driving the formation of a larger latent reservoir, or a beneficial effect, if the proviruses are driven into a permanently latent state, as proposed in the block-and-lock functional cure strategy (*40, 77*).

## Conclusions

We have identified a link between the role of LEDGF/p75 in HIV-1 replication and HIV-1 asRNA production. Although HIV-1 asRNA transcripts are frequently produced at low levels during HIV-1 replication, their levels were elevated by perturbation of the IN-LEDGF/p75 interaction. Furthermore, we showed that accumulation of HIV-1 asRNA in cells where IN-LEDGF/p75 interactions are perturbed was associated with reduced HIV-1 sense RNA and Gag protein levels, which is consistent with studies linking HIV-1 asRNA and IN-LEDGF/p75 perturbation to HIV-1 latency. Further characterization of the asRNA transcripts suggested that loss of IN-LEDGF/p75 interactions altered asRNA splicing.

ALLINIs efficiently block viral replication and have recently entered clinical trials for treatment of patients with HIV-1. Although the impact of elevated asRNA transcripts may be subtle, and in the presence of antiretrovirals the number of newly infected cells will be small, the potential link to latency and the role HIV-1 latency plays as a barrier to HIV-1 cure make this phenomenon worthy of study to better understand the potential impact of these novel therapeutics on HIV-1 cure strategies.

## Acknowledgments

We thank the NIH AIDS Reagent Program for generously providing reagents used in this work.

## Author contributions

Conceptualization: PRT, SGS

Investigation: MPC, DM, RS, OBU, CW, MG

Data analysis: THV, XG, WMM, DM, RS, MPC, OBU, PRT, MG, NK

Provided reagents: EMP, MK, ANE, VA, RTB

Funding acquisition: SGS, ANE, MK, WMM, PRT

Project administration: SGS, PRT

Writing – original draft: PRT, DM, SGS

Writing – review & editing: SGS, PRT, WMM, KAK, RTB, MK, EMP, ANE

## Data and materials availability

Data are available in the main text or the supplementary materials. Raw sequencing data and related analysis will be deposited and available in GenBank.

## Supplementary Materials for

### Materials and Methods

#### Cell culture

Jurkat cells were obtained through the NIH HIV Reagent Program, Division of AIDS, NIAID, NIH: Jurkat (E6-1) Cells, ARP-177, contributed by ATCC (Arthur Weiss) (*78*). They were cultured in RPMI 1640 (Invitrogen) supplemented with 10% fetal bovine serum (FBS) at 37°C with 5% CO^2^. *PSIP1* knock-out (LKO) and *HDGFL2* knock-out (HKO) Jurkat cells were provided by Eric Poeschla (*79*-*80*). P-TEFb mutant Jurkat cells were provided by Ryan T. Behrens and Nathan M. Sherer (*81*). HEK293T cells were maintained in Gibco DMEM including 10% FBS and 2 mM L-glutamine at 37°C with 5% CO^2^. LKO and CPSF6 knock-out (CKO) HEK293T cells were previously described (*17, 79*). Human CD4^+^T lymphocytes were isolated from peripheral blood mononuclear cells (PBMCs) (STEMCELL Technologies Lot# 1609270160) by activating and expanding T cells using 60 U/mL rh IL-2, and 25 µL/mL ImmunoCult^TM^ Human CD3/CD28/CD2 T Cell Activator (STEMCELL Technologies) for 12 days. Twenty-four h prior to use, activated T cells were cultured in RPMI 1640 media supplemented with 10% FBS at 37°C with 5% CO^2^. HEK293T cells were maintained in Gibco DMEM including 10% FBS and 2 mM L-glutamine at 37°C with 5% CO^2^.

#### Infection using HIV-1 virus

All viruses were produced by transfecting HEK293T/17 cells with HIV-1 pNL4-3 or recombinant pNL4-3 (NL4-3 ΔVpr Δenv). Recombinant viruses (HIV-1) were pseudotyped with VSV-G glycoprotein. Tat deficient viruses were obtained through the NIH AIDS Reagent Program, Division of AIDS, NIAID, NIH: Strain HXB2 ΔTat Non-Infectious Molecular Clone (pMtat(-)), ARP-2085 (*61*), and Strain HXB2 ΔTat Non-Infectious Molecular Clone (pMtat30), ARP-2087 (*62, 63*), were contributed by Reza Sadaie. Tat-deficient viruses were generated by co-transfection with Tat-expression vector under a CMV promoter. Tat-expression vector was obtained through the NIH HIV Reagent Program, Division of AIDS, NIAID, NIH: HIV-1 SF-2 Tat Eukaryotic Expression Plasmid (pCEP4-Tat), ARP-4691, were contributed by Lung-Ji Chang (*82*). Cells to be infected were seeded at a density of 7x10^4^ per well of a 24-well cell culture plate. ALLINI-treated cells were in media containing either 10 μM BI-D or BI-224436, 12 h prior to infection. Infections were done at a multiplicity of infection (MOI) between 0.1-1 and cells were washed 8, 24, and 36 hpi with 1x PBS and resuspended in fresh media. Cells were collected 48 hpi, unless otherwise stated. The number of infected cells were determined by vRNA expression, integrated vDNA, and/or p24 staining.

#### *In situ* vRNA detection

HIV-1 RNA in cells was detected using RNAscope^TM^ probing reagents (Advanced Cell Diagnostics). The manufacturer’s protocol was used with some modifications. Following fixation, cells can be stored in DPBS at 4°C for up to one week. Prior to hybridization, cells were washed with 0.1% Tween in PBS (PBST) for 10 min, and twice more in PBS for 1 min. Coverslips were immobilized on glass slides by placing a small drop of nail polish on a glass slide and the coverslip edge was placed on the nail polish drop. Sample was submerged in PBS to prevent dehydration. To prevent loss of reagent, a hydrophobic barrier was made by using an ImmEdge hydrophobic barrier pen (Vector Laboratories). The protease solution (Pretreat 3) as part of the manufacturer’s kit was diluted 1:15 in PBS and incubated in a humidified HybEZ oven (ACDBio) at 40°C for 15 mi. Nuclease treated samples were incubated with 0.25 μg or 0.5 units of DNase I in 1x DNase I Reaction Buffer at 37°C for 30 min or 0.5 μg RNase A in D-PBS at 37°C for 30 min. Proprietary pre-designed probes were added to the coverslip: RNAscope^TM^ Probe-HIV-gagpol-sense (ACDBio; catalog #317701) and RNAscope^TM^ Probe-HIV-nongagpol (ACDBio; catalog #317711-C3). Probes were incubated with the sample in a humidified HybEZ oven at 40°C for 2 h. The probes were then washed away with the wash buffer provided within the kit followed by two additional was steps. All wash steps were performed on a rocking platform at room temperature for 2 min. The probes were visualized by hybridizing preamplifiers, amplifiers, and finally, fluorescent label as described in the manufacturer’s protocol. The labeled probe set Amp 4A-FL, Amp 4B-FL, or Amp 4C-FL was used. Nuclei were counterstained with Hoechst 33342 (Invitrogen) at a concentration of 2 μg/ml for 4 min at room temperature. The counterstain was washed away with PBS at room temperature for 2 min following two additional wash steps. Coverslip samples were then mounted on slides using Prolong Gold Antifade (Invitrogen). Samples were analyzed the following day.

#### Immunostaining

Immunostaining for proteins in samples treated for vRNA detection was performed after probing for RNA using the protocol provided in this work. Coverslips were blocked with 5% bovine serum albumen (BSA) in 0.1% PBS-Tween-20 (PBST) at room temperature for 30 min. CA was probed with anti-p24 diluted 1:2,000 in 5% BSA in 0.1% PBST and incubated overnight at 4°C. Samples were washed twice in PBST at room temperature for 10 minutes each cycle. Goat anti-mouse secondary antibody (Invitrogen Catalog # A32727) was used at 1:1,000 dilution and incubated at room temperature for 1 h. The samples were then washed twice in PBST at room temperature for 10 min each cycle. Following immunostaining, nuclei were counterstained with Hoechst 33342 (Invitrogen) at a concentration of 2 μg/ml for 4 min at room temperature. The counterstain was washed away with PBS at room temperature for 2 min following two additional wash steps. Coverslip samples were then mounted on slides using Prolong Gold Antifade (Invitrogen). Samples were analyzed the following day.

#### Alu qPCR to measure integrated vDNA

DNA was isolated from HEK293T, Jurkat, and primary CD4^+^T cells all seeded at the same density 12 hpi using the DNeasy Blood & Tissue Kit (Qiagen). Samples were treated with RNase A as per instructions provided by the manufacturer. The two-step PCR approach was used to quantify the relative amount of provirus in each sample. Primers are listed in table S1. First round PCR: The 20 μL reaction included 13.32 µL PCR-grade H^2^O, 2.4 μL MgCl^2^ (4 mM final concentration), 0.04 µL Primer L-M667 (100 nM final concentration), 0.12 µL of each Alu primer (300 nM final concentration), 2 µL LightCycler FastStart DNA Master, and 2 µL of genomic DNA sample (concentration varies). Samples were mixed and lightly centrifuged prior to loading into the thermocycler instrument under the following parameters: Initial 8-min DNA polymerase activation step at 95°C. A 12-cycle target amplification step was conducted at 95°C denature phase for 10 sec, 60°C annealing phase for 10 sec, and a 72°C extension phase for 180 sec. A final 40°C cooling phase for 30 seconds was carried out prior to samples being plunged to 4°C for temporary storage. Second round PCR: Samples from first round PCR were diluted 10-fold prior to second round PCR. The 20 μL reaction included 12.96 µL PCR-grade H^2^O, 2.4 μL MgCl^2^ (4 mM final concentration), 0.12 µL of primers AA55M and Lambda T (300 nM final concentration), 2 µL LightCycler FastStart DNA Master, 0.2 µL of each Fluorogenic hybridization probe LTR FL and Fluorogenic hybridization probe LTR LC (200 nM final concentration), and 2 µL of diluted sample from the first-round PCR amplification. Samples were mixed and lightly centrifuged prior to loading into the thermocycler instrument under the following parameters: Initial 8-minute DNA polymerase activation step at 95°C. A 40-cycle target amplification step was conducted at 95°C denature phase for 10 sec, 60°C annealing phase for 10 sec, and a 72°C extension phase for 9 sec. Fluorescent measurement was carried out following the annealing step. A final 40°C cooling phase for 30 sec was carried out prior to samples being plunged to 4°C for temporary storage.

#### Quantitation and localization of fluorescence intensity for sense RNA, asRNA, and CA

HEK293T, Jurkat, and primary CD4^+^T cells were infected with HIV-1, VSV-G-pseudotyped virus, or delta-Tat variants at an MOI of 0.1-1. Cells were collected and washed three times with DPBS. After the final wash step, the cells were resuspended in 50 µL DPBS and seeded on poly-d-lysine coated coverslips for 30 min at a density to allow cells to be at least 1-2 cell diameters apart. Cells were immediately fixed with 4% PFA for 30 min. A Cytation 5 Cell Imaging Multimode Reader (BioTek) was utilized to measure fluorescence intensity within individual cells. To capture >500 infected cells containing sense RNA, asRNA, or CA at least five fields of view were quantified. Capture settings were not changed between fields-of-view or samples. For all fluorescence analyses, an uninfected negative control was used to measure background levels and only cells exceeding background were quantified. High content screening (HCS) CellMask™ Deep Red (Thermo Fisher Scientific) at 1:5,000 dilution and Hoechst 33342 (Invitrogen) at 1:5,000 dilution was used to mark cytoplasmic and nuclear compartments respectively for measuring asRNA levels.

#### Two-step RT-qPCR for sense and asRNA levels

RNA from HIV-1-infected WT and LKO Jurkat cells was isolated using the PureLink™ RNA Mini Kit (Thermo Fisher Scientific) with on-column DNase I treatment. Reverse Transcriptase (RT) reaction with Maxima H Minus RT was carried out using the manufacturer’s recommended protocol. The resulting cDNA was PCR amplified and detected following the LightCycler® 480 SYBR Green I Master protocol on a LightCycler 480 Instrument II from Roche. Specifically, for the asRNA transcripts, we utilized primers that amplify a region at the 5’ end of the molecules, for this region has been shown to be present in all asRNAs reported (*51*). Viral RNA from different sample preparations were normalized to GAPDH to derive relative quantities of sense and asRNA. The cycling parameters of quantitative PCR were as follows: (i) 2 min at 95°C; (ii) 40 cycles at 95°C for 15 sec, 60°C for 30 sec, acquisition. Strand-specific primer sequences can be found in table S1.

#### Library preparation for Illumina HiSeq

Activated human primary CD4^+^T cells (STEMCELL Technologies) were seeded in a 6-well plate and treated with 10 µM BI-D or equivalent volume of 100% DMSO as a vehicle control. Twelve h post treatment, cells were infected with HIV-1 (NL4-3) at an MOI of 1. Cells were harvested 24 hpi, and RNA extracted as a bulk population using Qiagen RNeasy Mini Kit. This process was performed three times to produce three independent data sets, each containing a DMSO control and a BI-D-treated sample. Ribosomal-RNA-depleted cellular RNA was generated using the Illumina Ribo-Zero Plus rRNA Depletion Kit, followed by the synthesis of stranded cDNA libraries using SMARTer Stranded RNA-seq (Takara Bio). The resulting libraries were sequenced at the Emory Non-human primate genomics core using the Illumina HiSeq approach and the resulting reads were aligned to the HIV-1 genome (NL4-3), using BLAT to assign position and sense.

#### Statistical analyses

All experiments were performed at least 3 times unless otherwise stated. Analysis and plotting of data were performed using the GraphPad Prism 7.0 software (GraphPad Software, La Jolla California USA) and are expressed as the mean ± SD. Statistical analyses were performed with Student’s *t*-test or One-Way ANOVA with Tukey’s multiple comparison test. Significance tests for plots analyzing percentage of vRNA expressing cells compared antisense RNA cells only (i.e. (-)RNA and (+/-)RNA expressing cells). Significant difference is defined as *p* < 0.05.

**Fig. S1.**
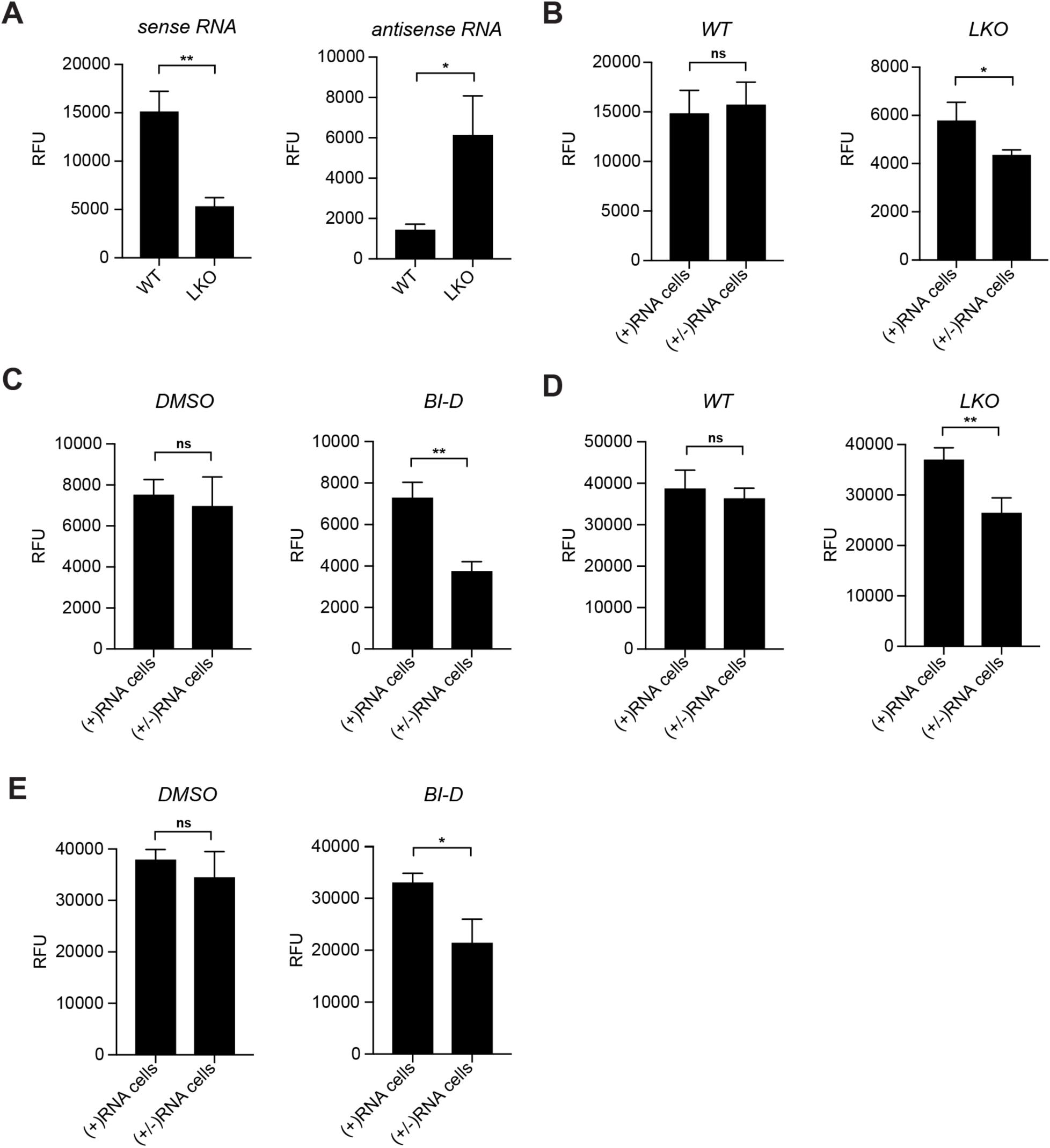
HIV-1 sense RNA and gag protein expression in the presence of asRNA. HEK293 cells were infected with VSV-G-pseudotyped virus, fixed, and labeled for vRNA and p24. Following image acquisition, mean fluorescence intensity was measured using by BioTek Gen5 software. (**A**) Mean fluorescence intensity of sense RNA (*left*) and asRNA (*right*) was plotted for WT and LKO HEK293 cells. (**B**) HIV-1 sense RNA levels in WT (*left*) and LKO (*right*) HEK293 cells containing either exclusively sense RNA ((+)RNA) or both vRNA transcripts ((+/-)RNA). (**C**) HIV-1 sense RNA level in DMSO-treated (*left*) and BI-D-treated (*right*) HEK293 cells. Cells were divided into two populations, those expressing sense RNA ((+)RNA) only, and those expressing both vRNA transcripts ((+/-)RNA). (**D**) HIV-1 gag protein levels in WT (*left*) and LKO (*right*) HEK293 cells, following separation into cells expressing (+)RNA only, and those expressing (+/-)RNA. (**E**) HIV-1 p24 levels in DMSO-(*left*) and BI-D-treated (*right*) HEK293 cells, following separation into cells expressing (+)RNA and those expressing (+/-)RNA. For all scoring conditions *n* = 3 independent experiments. At least 5 fields of view containing a total of >500 vRNA expressing cells scored for each experiment, with standard deviations shown. * p ≤ 0.05, ** p ≤ 0.01.

**Fig. S2.**
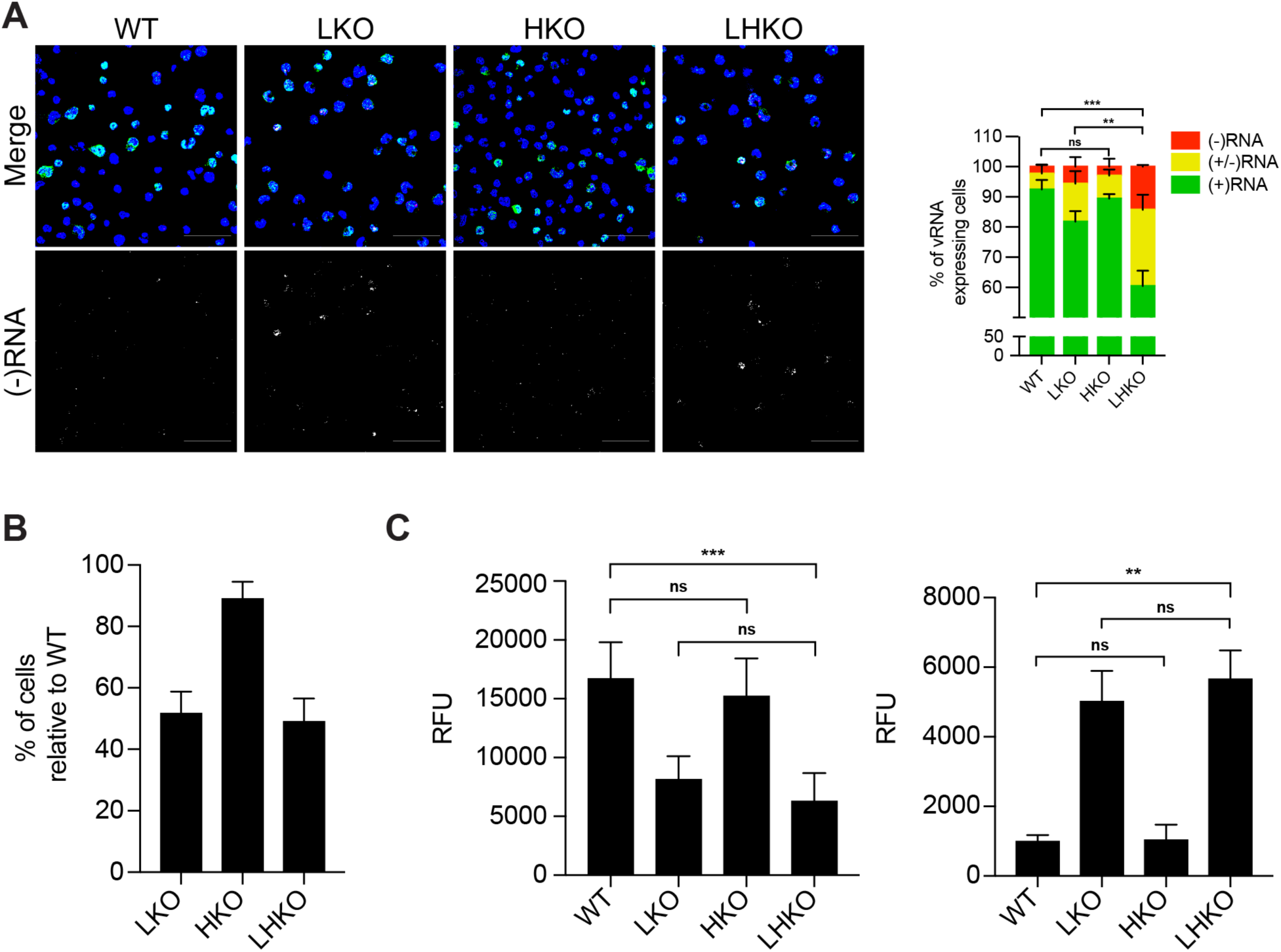
HIV-1 asRNA abundance in LEDGF/ HDGFL2 double knock-out and LKO cells. (**A**) RNA *in situ* hybridization was used to label HIV-1 sense RNA (green) and asRNA (red) in WT, LKO, HDGFL2 knock-out (HKO), and LEDGF/ HDGFL2 double knock-out (LHKO) Jurkat cells. The proportion of cells expressing sense RNA ((+)RNA), asRNA ((-)RNA), or both ((+/-)RNA) was plotted. (**B**) Number of cells producing vRNA was used to calculate infectivity of LKO, HKO, and LHKO Jurkat cell lines, relative to WT. (**C**) Mean fluorescence intensity of labeled HIV-1 sense (*left*) and antisense RNA (*right*) was measured in WT, LKO, HKO, and LHKO Jurkat cells. For all scoring conditions *n* = 3 independent experiments. At least 5 fields of view containing a total of >500 vRNA expressing cells scored for each experiment, with standard deviations shown. Scale bars represent 80 µm. * p ≤ 0.05, ** p ≤ 0.01, *** p ≤ 0.001.

**Fig. S3.**
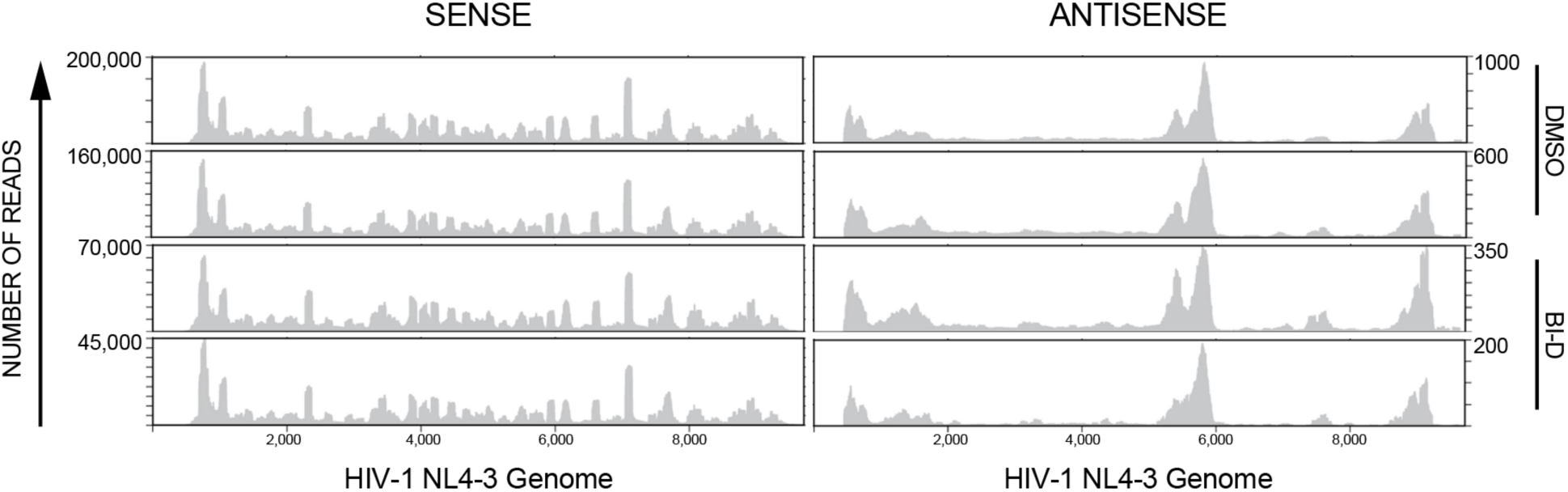
HIV-1 sense and antisense transcripts mapped to the genome. Activated primary CD4^+^ T cells were infected with HIV-1 (NL4-3) and treated with DMSO or an ALLINI to block IN-LEDGF/p75 interactions. After 24 hours, cells were harvested, and the RNA extracted as a bulk population. Stranded cDNA libraries were generated following ribosomal-RNA depletion. These libraries were sequenced using the Illumina HiSeq approach and the resulting reads were aligned to the HIV-1 genome, with position and sense assigned. Biological triplicates (two shown above, one in Fig. S4A) for each condition were sequenced.

**Fig. S4.**
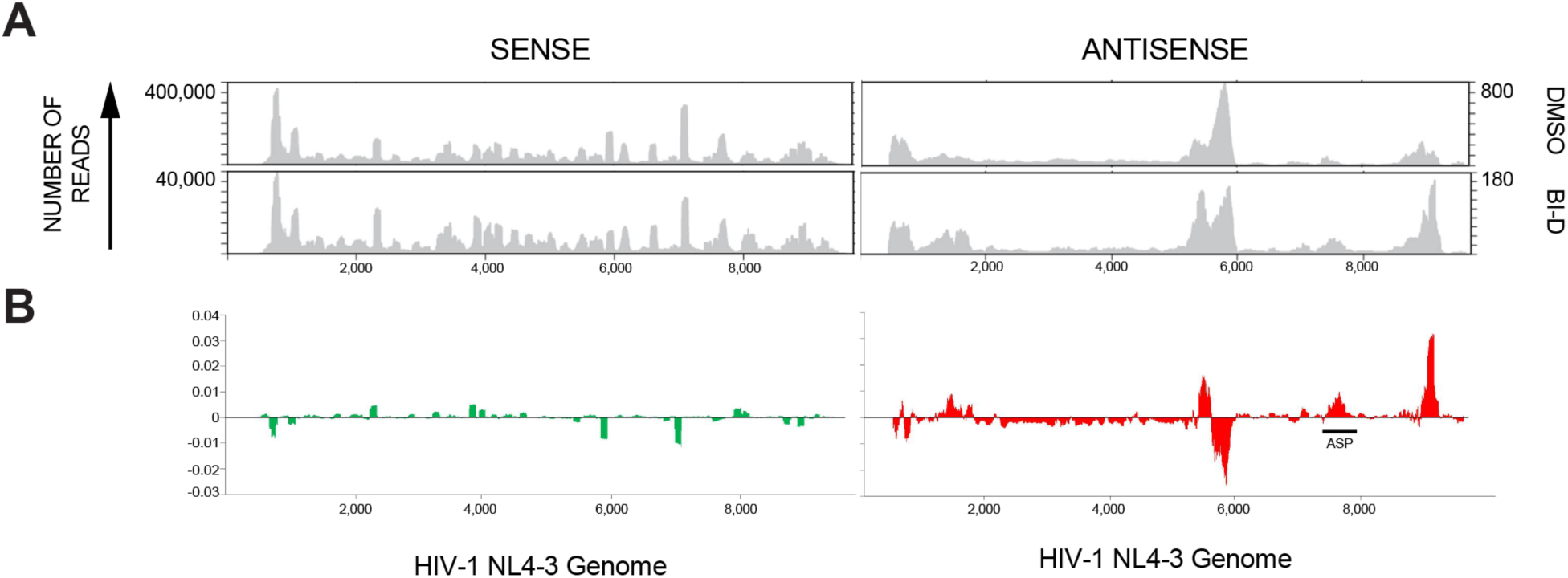
Characterization of HIV-1 asRNA transcripts by Illumina sequencing. (**A**) Activated primary CD4^+^T cells were infected with HIV-1 (NL4-3) and treated with DMSO or an ALLINI. After 24 h, cells were harvested and total RNA extracted; stranded cDNA libraries were generated following ribosomal-RNA-depletion. These libraries were sequenced using Illumina HiSeq and the resulting reads were aligned to the HIV-1 genome. Biological triplicates for each condition were sequenced; thus, three DMSO- and BI-D-treated samples were prepared. Representative read distribution plots are shown. (**B**) The reads at each position for each RNAseq plot were expressed as a proportion of all the aligned reads in the respective samples. The values for DMSO-treated cells were subtracted from those of ALLINI-treated samples to visualize changes in transcription following ALLINI treatment.

**Table S1.**
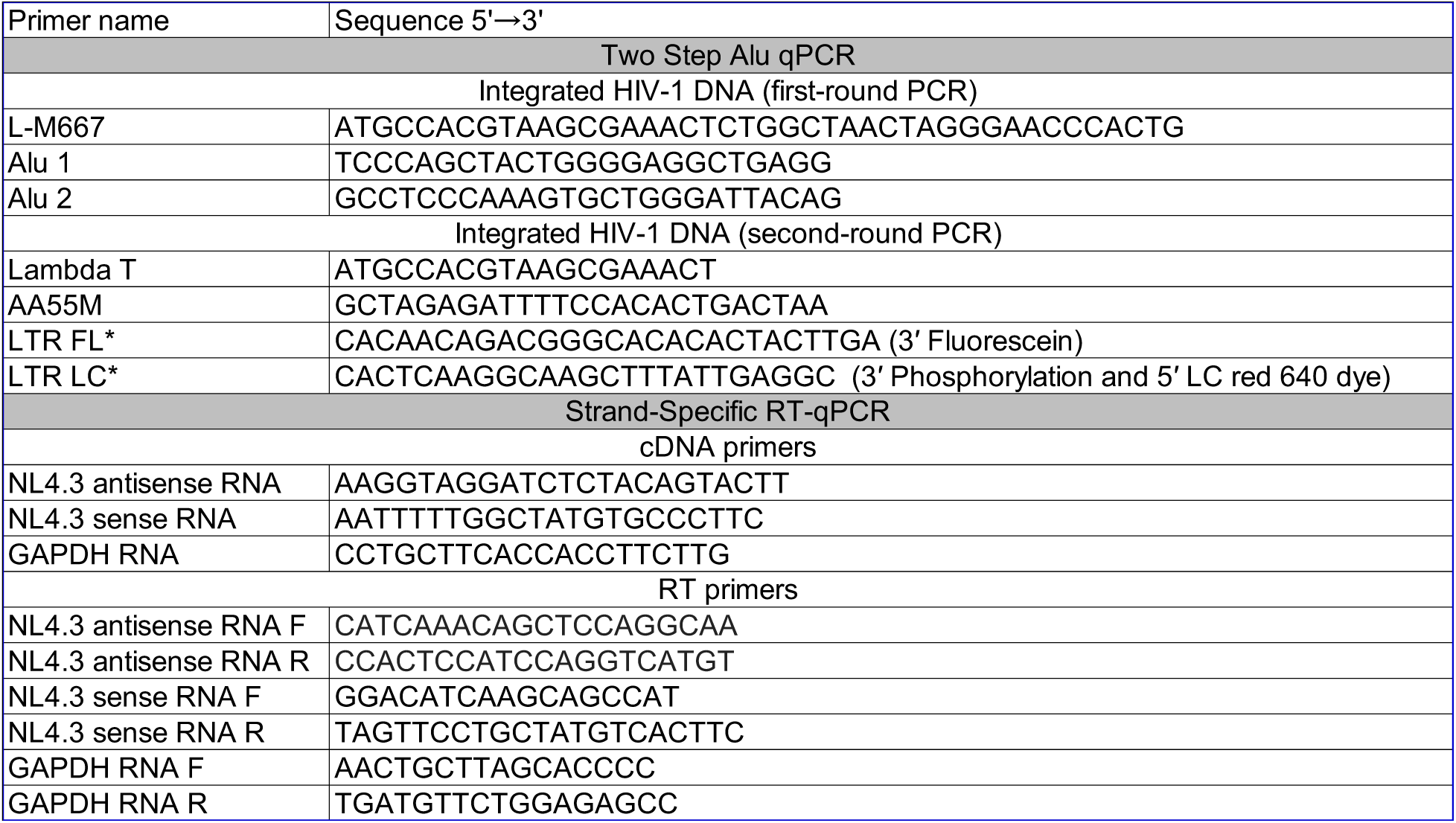
Primers used for two-step Alu qPCR and for strand-specific RT-qPCR for sense and asRNA determination.

